# Impact of single-cell RNA reference selection for the deconvolution of breast cancer spatial transcriptomics datasets

**DOI:** 10.64898/2026.01.09.698566

**Authors:** Stefan Altendorfer, Scott J. Walker, Carsten O. Daub

## Abstract

**Background:** Spot-based Spatial transcriptomics (ST) allows for unbiased gene expression analysis within tissue architecture, overcoming the limitations of single-cell RNA sequencing (scRNA-seq) by preserving spatial context. However, the high spatial resolution in ST leads to cellular heterogeneity within spots, requiring computational deconvolution to infer cellular compositions. While scRNA-seq serves as a key reference for deconvolution, the impact of reference composition on its accuracy is still unclear. In this study, we systematically evaluate the impact of reference selection for cellular deconvolution and give helpful guidelines to researchers to address this task.

**Methods:** We systematically evaluate the impact of single-cell reference selection for global and cell type-specific deconvolution performances in primary and metastatic breast cancer ST datasets. Focusing on state-of-the-art deconvolution tools Cell2location and Robust Cell Type Decomposition (RCTD), we assess the influence of varying reference sizes, cell-type distributions, reference-ST pairings, as well as the usage of large breast cancer and cross-cancer atlases.

**Results:** Our findings demonstrate that even small reference datasets can yield accurate deconvolution results, with RCTD and Cell2location exhibiting similar performances across cell-types. Heterogeneous entities, such as myeloids, presented greater challenges for accurate deconvolution compared to more distinct ones, such as epithelial cells. Spatial domains associated with prominent cell types like cancer and stromal cells were detected, although their contributions were systematically under- or over-estimated Additionally, we found that perfect reference-ST matching enhances deconvolution accuracy compared to the usage of cross-patient references. Finally, large breast cancer single-cell atlases composed of a diverse set of patients were also able to provide reliable deconvolution results.

**Conclusion:** This study provides important insights into optimizing reference selection for ST deconvolution, highlighting the strengths and limitations of current computational tools in addressing cellular heterogeneity within spatial transcriptomics datasets.

## 1 Background

Breast cancer remains the most prevalent malignancy among women worldwide, accounting for over 2.3 million new cases and approximately 670,000 deaths annually[1]. Despite advances in early detection and treatment, metastatic breast cancer continues to pose significant clinical challenges, contributing to the majority of breast cancer-related mortality[2, 3]. The disease exhibits profound intra- and inter-tumoral heterogeneity, with molecular subtypes defined by the expression or absence of the hormonal receptors estrogen receptor (ER), progesterone receptor (PR), and human epidermal growth factor receptor 2 (HER2) into Luminal A/B, HER2-enriched, and Triple-Negative Breast Cancer (TNBC), which in turn guide therapeutic decisions[4–7]. However, these broad classifications fail to capture the full complexity of tumor biology and its dynamic, heterogeneous microenvironment.

Single-cell RNA sequencing (scRNA-seq) has revolutionized cancer biology by enabling high-resolution profiling of cellular heterogeneity within tumors and their surrounding microenvironments[8]. In breast cancer, scRNA-seq has aided in the elucidation of the roles of cancer-associated fibroblasts (CAFs), tumor-infiltrating lymphocytes (TILs), and myeloid subsets in tumor progression, immune evasion, and therapeutic resistance[9, 10]. Nevertheless, scRNA-seq inherently lacks spatial context, limiting its ability to resolve cell-cell interactions and tissue architecture.

Spatial transcriptomics (ST) overcomes this limitation by profiling gene expression within intact tissue sections while preserving spatial context[11, 12]. Among the various ST technologies, spot-based platforms such as 10x Genomics Visium have gained wide adoption in breast cancer research. These approaches capture transcripts across spots with diameters of 55 *µ*m, each typically containing multiple cells. While this enables unbiased transcriptome-wide profiling, each spot becomes a miniature bulk RNA sample complicating downstream biological interpretation at a cellular level. [13] In addition to spot-based approaches, fluorescence-imaging-based spatial transcriptomics methods such as MERFISH[14] and 10x Genomics Xenium[15] capture RNA transcripts at single-cell, or sub single-cell resolution, although they are limited by targeted gene panels[16].

To address the heterogeneity within spatial spots a wide range of computational methods have been developed to perform cellular deconvolution. These tools infer the relative abundance of different cell types contained within each sequenced spot by leveraging scRNA-seq reference dataset(s), allowing for finer-grained insights into tumor architecture and cell-cell communication. [17] In particular, the tools Cell2location[18] and RCTD[19] have demonstrated strong overall performance in recent benchmarking studies[20–23].

Despite advancements in this analysis, one important factor has received little attention, the choice of the scRNA-seq reference dataset. Most benchmarks rely on single references, overlooking the impact of reference size, cell-type diversity, and patient matching which may substantially influence deconvolution outcomes, with consequences for biological interpretation and reproducibility.

In this study, we systematically evaluate how scRNA-seq reference selection impacts spot-based ST deconvolution in ductal carcinoma in situ (DCIS) and metastatic breast cancer (MBC). Specifically, we investigate the influence of reference cell numbers, relative abundance of particular cell types, the role of patient and/or disease subtype matching, and the utility of multi-patient or cross-cancer reference panels. Our findings provide practical guidelines for researchers designing ST studies, evaluated the importance of reference selection when integrating scRNA-seq and ST for breast cancer research, and give insights into the necessity of creating a matched scRNA-seq dataset along with ST experiments.

## 2 Methods

### 2.1 Datasets

We analyzed breast cancer (BC) Xenium and MERFISH in situ spatial transcriptomic sequencing data. Single-cell RNA-seq (scRNA-seq) datasets were used as reference panels for cell deconvolution, whereas two Xenium and three MERFISH samples served as targets for inferring cell-type proportions in spatial contexts. All datasets used in this study are publicly available. References, sources and further metadata for each dataset can be found in Supplementary Tables **1 and 2**.

**Fig. 1.**
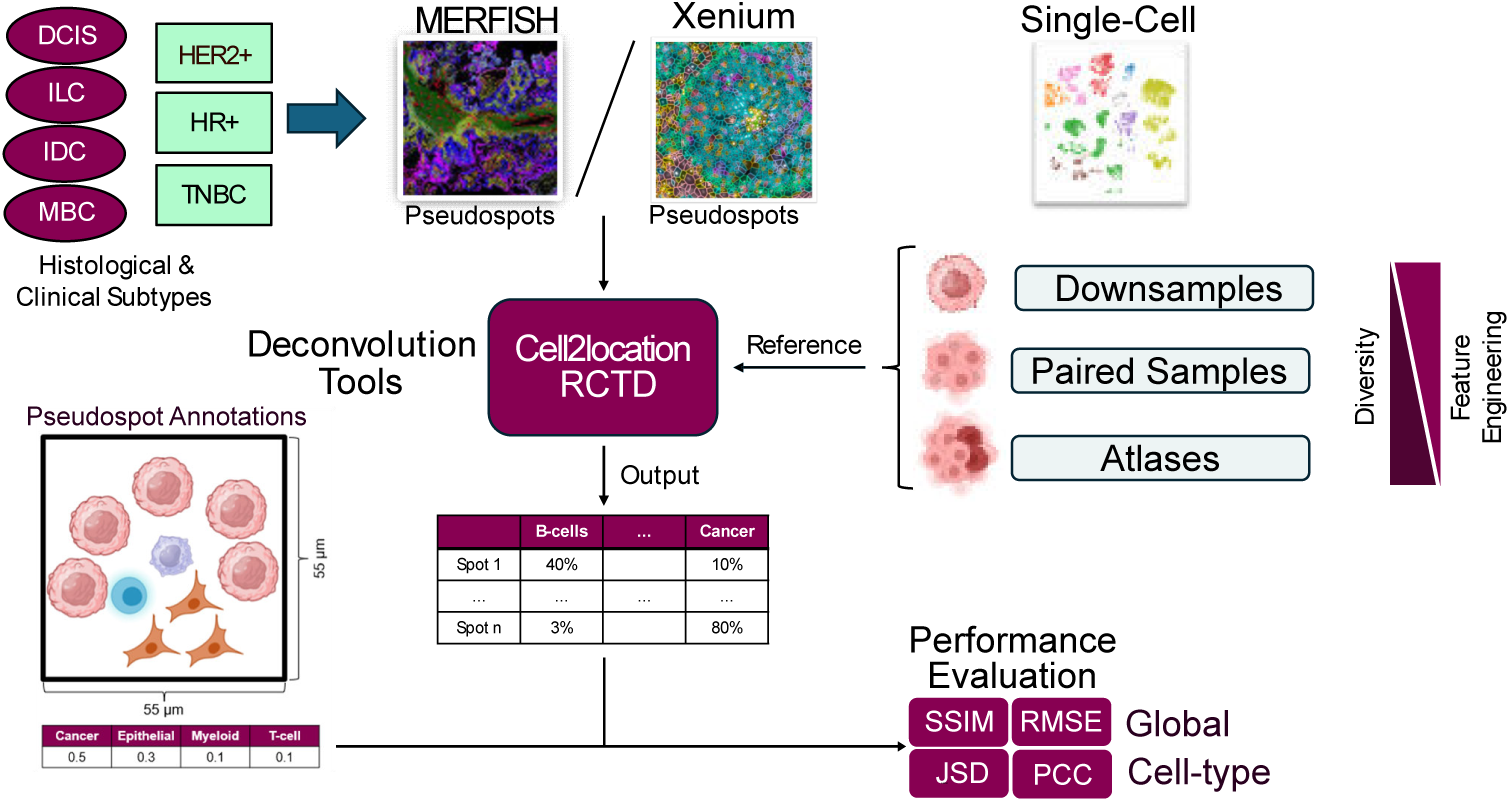
Study overview for investigating the impact of scRNA-seq reference selection for cellular deconvolution of breast cancer ST datasets. Parts of the figure were created with BioRender and technology images adapted from https://accela.eu/products/merscope-platform-vizgen/ and https://www.10xgenomics.com/support/software/xenium-explorer/latest.

### 2.2 Data preprocessing and quality control

#### scRNA-seq datasets

All scRNA-seq datasets were collected using 10x Genomics Chromium protocol. Raw count matrices from `CellRanger` analysis software[24] were analyzed in Python using the `Scanpy`[25] tool. Cells with fewer than 200 unique molecular counts, and genes expressed in fewer than three cells were removed. Doublets, where two cells are captured and sequenced together with the same barcode, were predicted and removed using the `Scrublet`[26] tool. Cell-type annotations were provided by the original authors. These were curated into harmonized major and minor categories.

#### Xenium and MERFISH datasets

Both Xenium and MERFISH samples were segmented by Janesick `et al.`[15] and Klughammer `et al.`[27] respectively. Downloaded Xenium bundles were processed using Python with the `spatialdata-io` tool[28]; Raw count matrices were used. Quality control included filtering cells with less than 10 transcript counts, and genes represented in fewer than 5 cells. Expert-curated annotations from the original authors were used for cell-type labeling, harmonized to allow annotation matching to the single-cell references and ambiguous hybrid cells were removed.

### 2.3 Pseudospot creation for benchmarking

Cell deconvolution is performed on spot based spatial transcriptomics datasets primarily, e.g. 10x Genomics Visium protocol. To mimic Visium spot resolution, segmented Xenium and MERFISH cells were aggregated into pseudospots by binning within a 55 *µ*m square grid. Gene expression profiles from all cells within each pseudospot were summed while preserving cell-type composition. These pseudospots provided ground truth proportions for benchmarking deconvolution.

### 2.4 Reference downsampling

To evaluate the impact of reference panel size and composition, we applied two downsampling strategies to the scRNA-seq datasets. In the stratified downsampling approach, we sampled random subsets of cells while preserving the original cell-type proportions. In the equal allocation downsampling approach, we instead sampled equal numbers of cells from each cell type to enforce balanced representation. Each down-sampling level was repeated with multiple random seeds to ensure robustness and to remove unintentional biases from cell selection.

### 2.5 Deconvolution algorithms

We utilized two widely used deconvolution tools. Cell2location[18] is a Bayesian hierarchical model estimating absolute cell-type abundances using negative binomial regression. This tool utilizes an underlying variational autoencoder from scVI[29]. RCTD[19] is a Poisson-based regression model projecting scRNA-seq-derived expression profiles onto spatial spots, allowing mixtures of multiple cell types per spot. Both methods were implemented following official documentation, with minor parameter adjustments for breast cancer datasets.

### 2.6 Performance metrics

Predicted cell-type proportions were compared against known ground truth values from pseudospots.

#### Global metrics

Global metrics summarize the overall accuracy of the model across all spots, without distinguishing between different cell types. We used root mean squared error (RMSE) as a point-wise error metric, penalizing large deviations between predicted and true proportions and ranging from 0 (no deviations) to infinity. Additionally, we leveraged structural similarity index (SSIM), which evaluates global agreement in variance and covariance between predicted and observed proportion matrices, ranging from -1 (perfect negative correlation) to 1 (perfect positive correlation).

#### Cell-type-specific metrics

To quantify performance for each individual cell type, we computed Pearson correlation coefficients (PCC) between predicted and true proportions, capturing linear agreement and ranging from -1 (perfect negative correlation) to 1 (perfect positive correlation). We also assessed Jensen–Shannon divergence (JSD), which measures the similarity between predicted and true distributions. JSD values range from 0 to 1, with lower values reflecting greater similarity. In addition, we calculated root mean squared error (RMSE) for each cell-type individually.

#### Meta-cell creation

Meta-cells where created using SEACells[30]. The recommended number of 72 cells per meta-cell were chosen. The meta-cells were constructed on the scVI[29] latent embedding matrix rather than the default of PCA. Meta-cells were checked to ensure the value of ‘celltype-purity’ was 80%.

## 3 Results

### 3.1 How does the number of reference cells affect deconvolution performances?

Single-cell RNA sequencing (scRNA-seq) datasets vary widely in size, with each cell representing a distinct experimental unit. Larger datasets offer improved statistical power and finer resolution of cellular heterogeneity, enabling more accurate identification of cell types, subpopulations, and rare states. [10, 31] However, the minimum number of cells required for effective cell deconvolution in spot-based spatial transcriptomics (ST) remains unclear. To address this, we systematically investigated how reference size and cell-type distribution affect deconvolution performance.

We applied two complementary downsampling strategies to scRNA-seq reference datasets: stratified downsampling, which preserves the original cell-type proportions, and equal allocation downsampling, which enforces balanced representation across cell types.

To benchmark deconvolution accuracy, we generated pseudospots *in silico* from segmented Xenium replicates provided by Janesick `et al.`[15] that represented a HER2+/HR+ ductal carcinoma in situ (DCIS) breast cancer subtype. This method allowed us to simulate 50 *µm* spots that could be collected from imaging-based ST protocols whilst also maintaining ground truth cell-type proportions.

All downsampling experiments were performed using matched scRNA-seq references derived from the same tumor and patient as the corresponding ST sample, thus limiting biologically derived technical batch effects. For the Xenium dataset, we employed two annotation levels: a major level comprising eight broad cell types, and a minor level with 17 subtypes, including distinct immune and stromal populations that are relevant to the breast cancer tumor microenvironment.

### 3.2 How many cells in total are required for deconvolution?

First, we examined how the total number of reference cells affected deconvolution by generating panels of 500–1,400 cells through stratified downsampling from the matching single-cell reference.

At a reference size of 1,400 cells, RCTD and Cell2location demonstrated robust performances, accurately reconstructing the spatial domains of cancer epithelial cells, cancer-associated fibroblasts, and T cells. Cancer cells were correctly localized to tumor-rich regions, whereas stromal cells populated adjacent tissue zones. T cells, although sparse in ductal carcinoma in situ (DCIS)[32], were moderately detected in stromal regions (refer **Fig. 2a**). Despite promising results, residual analysis revealed systematic biases. Fibroblasts were slightly underestimated in the stromal regions, whereas cancer cells were frequently overestimated by RCTD in the surrounding tumor domains (refer **Fig. 2b**). Quantitative evaluation using the Pearson correlation coefficient (PCC), Jensen-Shannon divergence (JSD), and root mean squared error (RMSE) confirmed the trends observed in the RCTD deconvolution (refer **Fig. 2c**). Cancer cells exhibited high PCC (0.96), low JSD (0.05), and reasonably low RMSE (0.13) values, indicating a strong agreement between the predicted and true proportions. Stromal cells showed high PCC (0.91) and low JSD (0.02) but a slightly higher RMSE (0.20) value, suggesting that the model captured the spatial distribution of fibroblasts but had problems in accurately quantifying them in a point-wise manner. T cell predictions were characterized by moderately high PCC (0.86), reasonably low JSD (0.14), and low RMSE (0.07) scores, reflecting an overall accurate representation. Interestingly, the tool Cell2location differed quite strongly in terms of systematic biases by slightly underestimating cancer cells in tumor domains and strongly overestimating T cells in stromal regions where cancer-associated fibroblasts were in turn severely underestimated (refer **Suppl. Fig. 1**).

**Fig. 2.**
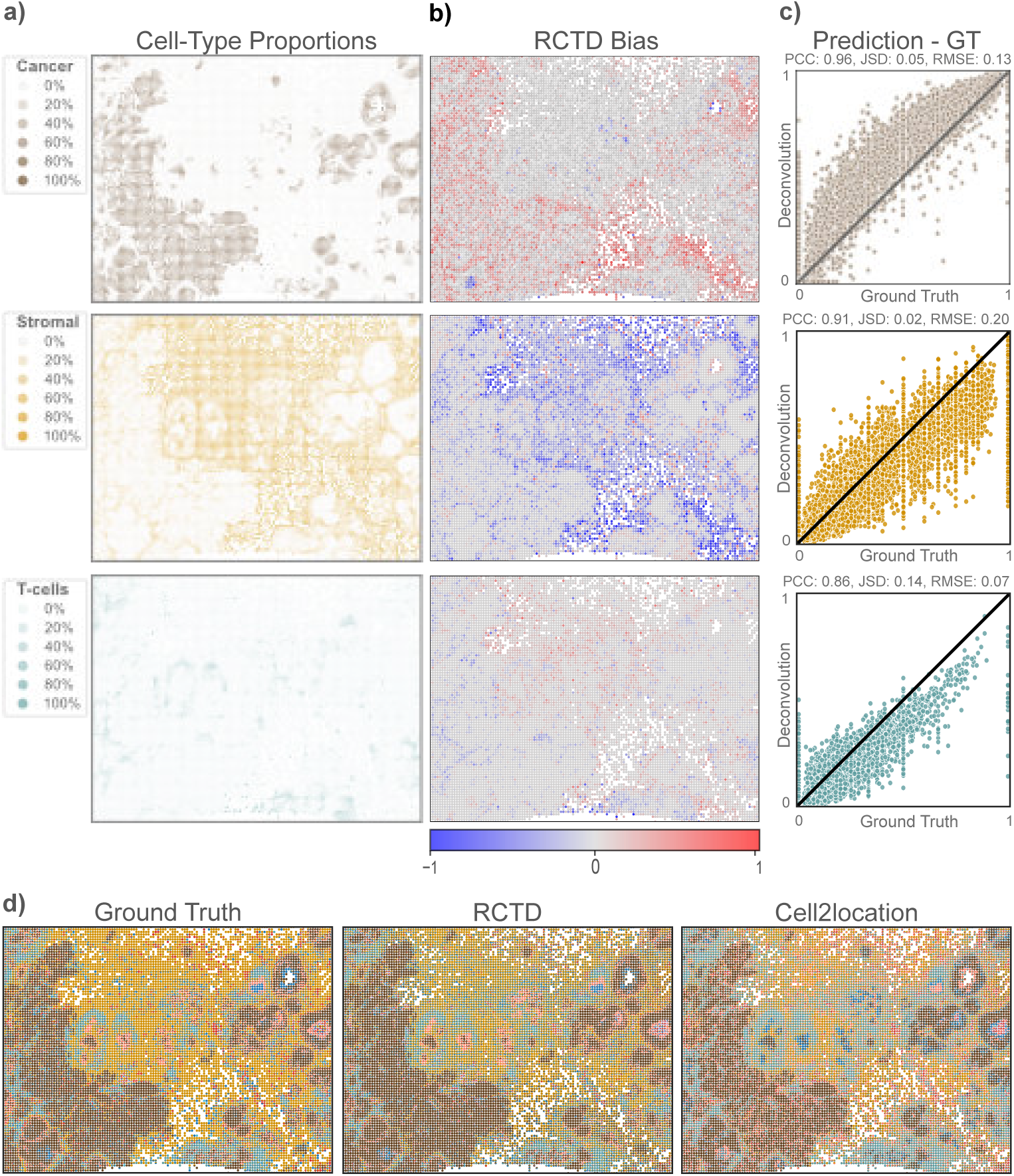
RCTD deconvolution analysis results for three key cell types: cancer cells, stromal cells, and T cells. **a)** Cell-type proportions per spatial spot based on segmented Xenium cell annotations. **b)** Difference between deconvolved proportions and ground truth for each spot, with overestimation indicated by red and underestimation by blue. **c)** Correlation plots comparing predicted and ground truth proportions for each spot, with the diagonal line representing perfect prediction. Performance metrics, including Pearson Correlation Coefficient (PCC), Jensen-Shannon Divergence (JSD), and Root Mean Squared Error (RMSE), are provided to quantify the accuracy of the predictions. **d)** Deconvolution results from RCTD and Cell2location by representing each spot as a pie chart, with each section indicating the proportion of different cell-types present in that spot.

To visualize the deconvolution results across cell types, we overlaid pie charts of cell-type proportions across spatial spots (refer **Fig. 2d**). The overall tissue architecture comprising large tumor domains, stromal regions, and immune cell clusters that appeared in the ground truth data were effectively captured by both deconvolution algorithms. Again, it became visible that Cell2location showed stronger deviations by overestimating immune cells across the ST slide. Correlation matrices of residuals revealed that the high abundance of one cell type distorted predictions for others, such as stromal cells affecting the predictions of immune cells. The most pronounced systematic biases arose when a singular cell type was highly abundant within spatial spots, leading to its own underestimation (refer **Suppl. Fig 4**).

To analyze how many cells of a matching single-cell reference are required for successful deconvolution, we evaluated performance across a gradient of downsampling levels. Cell2location and RCTD both maintained stable results (RCTD: RMSE *≈* 0.10, C2L: RMSE *≈* 0.12) down to 500 total cells, with no significant improvements in SSIM and RMSE (refer **Fig. 3a**) even compared to the full capacity of 20,000 cells (refer **Fig. 4a**). This illustrates that even small reference datasets are sufficient for accurate deconvolution. Notably, RCTD outperformed Cell2location in both metrics, consistent with previous results.

**Fig. 3.**
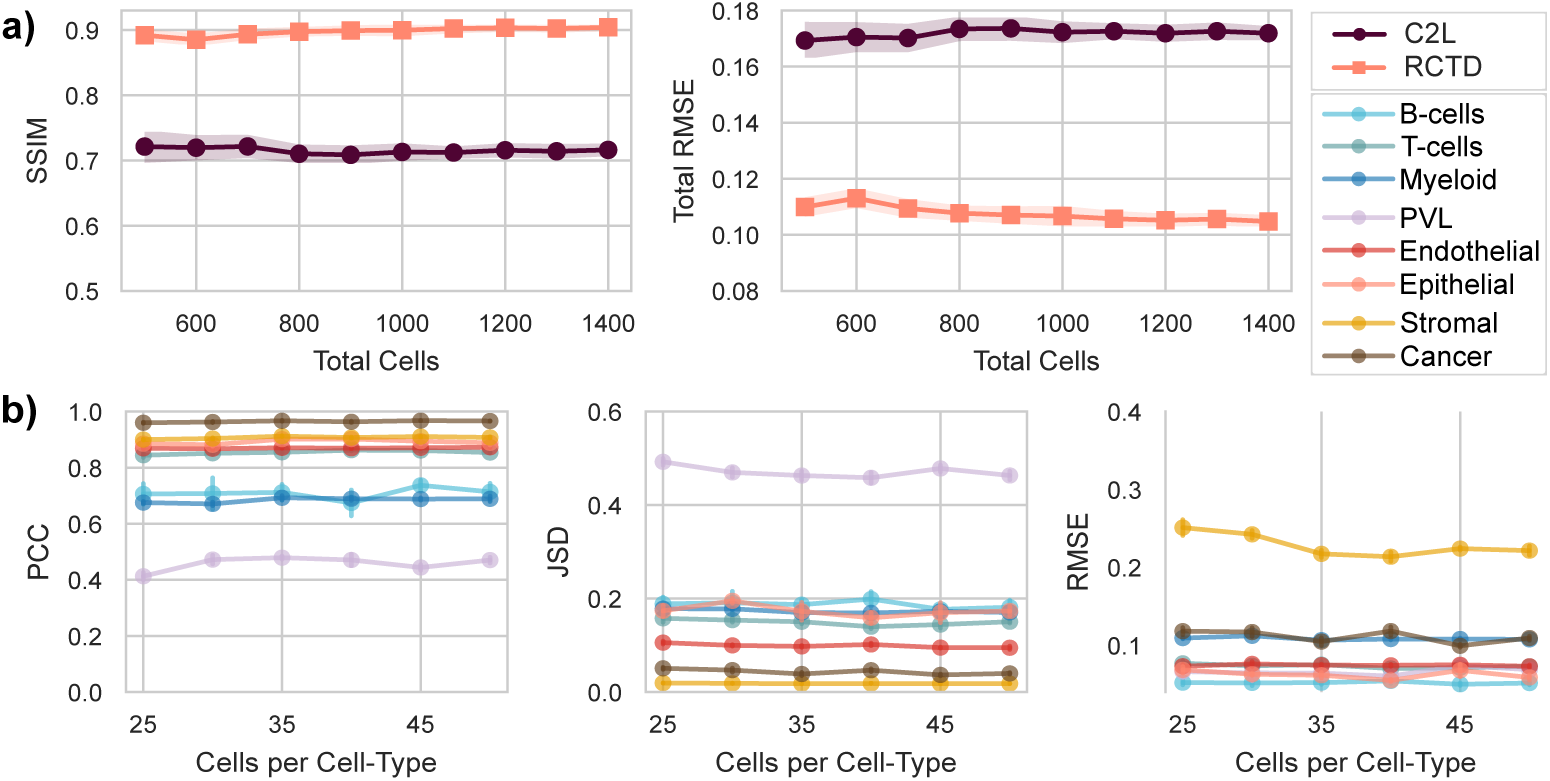
**a)** Global performance metrics RMSE (root mean squared error) and SSIM (structural similiarity index) across different downsampling levels for Cell2location and RCTD for the major annotation level in Xenium pseudospots. **b)** Equal allocation downsampling results of RCTD for every major cell-type. Deconvolution evaluation illustrated with cell-type specific evaluation metrics: PCC (pearson correlation coefficient), JSD (Jensen-Shannon divergenceJ and RMSE. Errorbars illustrate standard error (se) across deconvolution results of downsampling replicates.

**Fig. 4.**
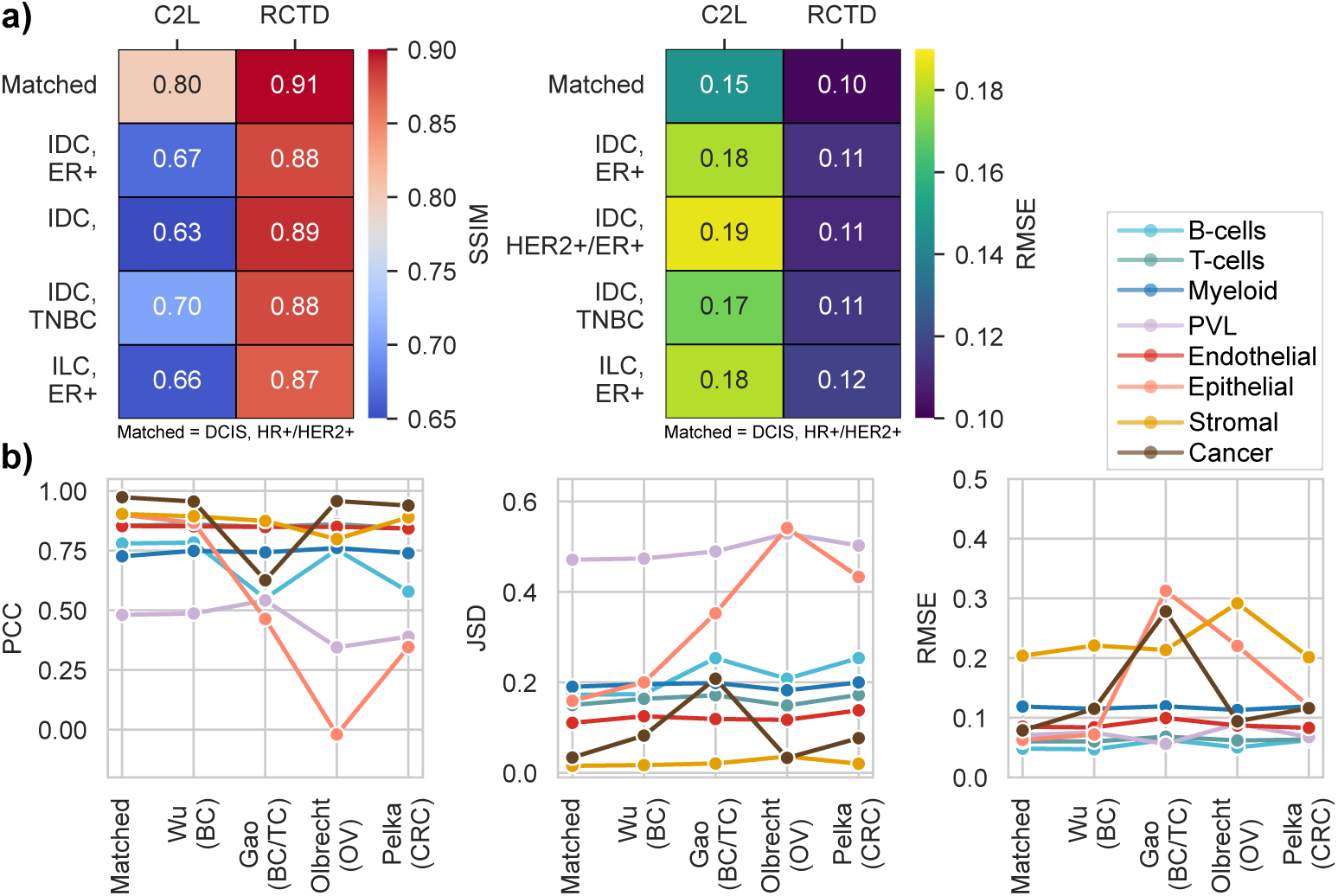
**a)** Global deconvolution performances of RCTD and Cell2location by using different single-cell samples with diverse clinical backgrounds as references. Shown values are the mean performances across both Xenium replicates 1 and 2. **b)** Cell-type-specific deconvolution performances of RCTD across different reference atlases from breast- or cross-cancer single-cell datasets.

#### 3.2.1 How many cells per cell-type are necessary?

Next, we evaluated the impact of cell-type distribution within scRNA-seq reference datasets for deconvolution and asked whether more heterogeneous cell types demand a higher representation in the reference dataset. We downsampled each cell type to equal numbers, with a minimum of 25 cells per type, which is the threshold required by RCTD. Overall, increasing the number of cells did not substantially improve deconvolution performances for either RCTD or Cell2location (refer **Fig. 3b** and **Suppl. Fig. 2**).

Cell-type specific performances varied widely across cell types. Cancer cells and cancer-associated fibroblasts (CAFs) achieved the highest spatial correlation. However, stromal cells were systematically underestimated, yielding a high RMSE (0.25). The least frequent cell type present in the tissue, perivascular-like cells, performed poorly in correlation-based metrics but showed low RMSE, suggesting stable but spatially uninformative predictions. Endothelial cells, epithelial cells, and T cells were consistently well captured across the metrics. Myeloid populations showed weaker performances likely because of their broad categorization. Additionally, cell-type rankings and evaluations were very similar to the results generated by the stratified downsampling approach (spearman correlation coefficients *>* 0.9 across all cell-type metrics), indicating that cell-type proportions within references has minimal impact on deconvolution results (refer **Suppl. Fig 2 and 3**).

To evaluate deconvolution success on a finer-grained annotation level, we leveraged 17 minor cell-type labels from the Janesick `et al.`[15] Xenium samples. This allowed us to further focus on cancer and immune cell heterogeneity. Interestingly, invasive tumor and in-situ subclusters showed divergent behaviors (refer **Suppl. Fig. 5a**). Invasive tumor predictions were more variable across replicates, while DCIS subtypes were reconstructed with greater accuracy by RCTD. Cell2location showed better performance when the broader ‘cancer’ annotation was used. In contrast, both algorithms more faithfully deconvolved CD4+ and CD8+ T cell populations than when using the broader “T cell” annotations (refer **Suppl. Fig 5b**). Further subtyping the broad category of myeloids also yielded better deconvolution performances (refer **Suppl. Fig. 5c**) since especially dendritic cells (DCs) and mast cells were quite distinguishable in their gene expression programs. These results indicated that finer-grained subtyping can improve resolution and deconvolution performances for certain cell-types. However, global RMSE and SSIM metrics, demonstrated an overall loss of accuracy due to the more complicated assignment task (refer **Suppl. Fig. 2**).

Overall, these results highlight that a relatively small number of reference cells is sufficient for many cell-types, but their corresponding subtyping level can also change deconvolution results drastically.

### 3.3 Are matched scRNA-seq references necessary for accurate deconvolution?

Given the large number of publicly available scRNA-seq datasets and the comparatively high cost of generating new reference data, it is critical to determine whether deconvolution algorithms benefit most from patient- and tumor-matched references, or whether references derived from other samples or breast cancer subtypes provide comparable performances. To address this question, we systematically quantified the effect of reference selection by comparing matched scRNA-seq samples with non-matching ones in Xenium pseudospots.

We evaluated deconvolution performances using the matched scRNA-seq dataset from Janesicket al.[15], representing a DCIS hormone receptor–positive (HR+) and HER2-positive (HER2+) subtype. This dataset was used as the control reference. In addition, we tested four non-matching references from Wu `et al.`[33] that encompassed distinct tumor subtypes, including HR+ invasive ductal carcinoma (IDC), HR+ invasive lobular carcinoma (ILC), and triple-negative breast cancer (TNBC).

Global metrics highlighted clear advantages of matched references when using Cell2location, which achieved its lowest RMSE at 0.15 and highest SSIM at 0.80 (refer **Fig. 4a**). By contrast, RCTD produced consistent results across reference pairings (SSIM = 0.88, RMSE = 0.11, with only slight improvement when using the matched reference (SSIM = 0.91, RMSE = 0.10).

Cell-type-specific evaluations further elucidated that epithelial predictions benefited most from matched references with Cell2location, while immune and stromal cell predictions showed negligible improvement (refer **Suppl. Fig. 6b**). RCTD on the other hand, achieved more stable cell-type-specific performances across reference-pairings (refer **Supp. Fig. 6a**). Visualizations of deconvolution outcomes (refer **Suppl. Fig. S7**) demonstrated that the inferred tissue architectures did not change depending on the reference used.

### 3.4 How stable is cell deconvolution to changes in sample heterogeneity?

Next, we assessed how increasing reference heterogeneity affects deconvolution performance by using large single-cell atlases as references for both Xenium replicates. We evaluated whether including more diverse sample cohorts improves or compromises prediction accuracy. Specifically, we compared the breast cancer atlas from Wu `et al.` 2021[33], the cross-cancer (breast and thyroid) atlas from Gao `et al.` 2021[34], and non–breast cancer atlases from ovarian (Olbrecht `et al.` 2021)[35] and colorectal cancer (Pelka `et al.` 2021)[36], benchmarked against the matched Janesick `et al.` 2021[15] scRNA-seq dataset as a gold-standard reference.

Global performance trends showed that both Cell2location and RCTD reached their highest accuracies when using the matched reference (RCTD: SSIM = 0.91; C2L: SSIM = 0.80; **Suppl. Fig. 8**). Using the heterogeneous Wu `et al.` breast cancer atlas[33] caused only modest performance drops (RCTD: SSIM = 0.89; C2L: SSIM = 0.70). The Gao cross-cancer dataset[34] yielded competitive results for Cell2location (SSIM = 0.75), but substantially reduced RCTD performance (SSIM = 0.68), largely due to misclassification of cancer cells. Non–breast cancer atlases had more pronounced effects: Cell2location’s performance decreased markedly (SSIM = 0.57), particularly in epithelial populations, whereas RCTD remained robust with the Pelka colorectal atlas (SSIM = 0.89) but showed lower accuracy with the ovarian reference (SSIM = 0.77).

Cell type–specific analysis (refer **Fig. 4b**) revealed that most non-cancer cell types were robustly deconvolved by both methods across all reference atlases, with only minor fluctuations for B cells when assessing their spatial distribution. Epithelial cells, however, were the major exception: accurate deconvolution was only achieved using breast cancer references, while performance declined sharply with non-breast atlases, highlighting the importance of tissue of origin for this cell type. In contrast, cancer cells were generally well recovered across different cancer datasets, suggesting that both Cell2location and RCTD are capable of picking up consensus cancer gene programs across tumor types. The only notable exception occurred with RCTD using the ovarian atlas, where it became spatially evident that the tool failed to distinguish cancer from normal epithelial cells, resulting in substantial overestimation of epithelial abundance within tumor domains (refer **Suppl. Fig. 9**).

Taken together, these results show that while matched references still remain optimal, large heterogeneous atlases of the same cancer type can perform comparably.

### 3.5 Expanding deconvolution to metastatic MERFISH datasets

In addition to the analyses conducted on the DCIS Xenium replicates, we also evaluated deconvolution performances on metastatic MERFISH liver samples from Klughammer `et al.` 2024[27]. This analysis was conducted to validate our findings across different ST protocols, as well as a different sample tissue. We chose liver metastatic samples as they contained plenty of subtype-matched single-cell references. However, MERFISH experiments contained only a fraction of spots (around 1,700) compared to the Xenium replicates (around 12,000) and consisted of only 7 major cell-types (B-cells, endothelial cells, stromal cells, hepatocytes, MBC cells, myeloids and T-cells).

Evaluating downsampling performance on matched MERFISH-single-cell pairs further demonstrated that increasing the number of cells did not substantially improve deconvolution results for neither RCTD (RMSE *≈* 0.09, SSIM *≈* 0.93) nor Cell2location (RMSE *≈* 0.11, SSIM *≈* 0.89) (refer **Suppl. Fig. 12**). The overall tissue architecture comprising cancer domains, hepatocyte clusters, and stromal tissue with infiltrating immune cells was effectively captured by both deconvolution tools (refer **Suppl. Fig. 11a**). However, both methods struggled to accurately resolve cancer cells at the individual level (refer **Suppl. Fig. 13**. Notably, Cell2location again tended to underestimate the cancer cell contribution in tumor-dense spots (refer **Suppl. Fig. 11b)**. B-cells, being the rarest cell type in the MERFISH sample, presented a challenge for deconvolution with both tools failing to capture their spatial distribution indicated by low PCC (0.53) and high RMSE (0.41) values. However, neither systematic overestimation nor underestimation was observed (RMSE = 0.08), similar to what was seen with PVL cells in the Xenium replicates. Moderately abundant cell types (in this case endothelial cells, fibroblasts, and myeloid cells) exhibited again the best performances across cell-type-specific metrics (RMSE *<* 0.10, PCC *>* 0.84 and JSD *<* 0.14).

To further assess the impact of correct subtype matching, we aggregated the metastatic breast cancer scRNA-seq samples from Klughammer `et al.` 2024[27]. These included HER2+, HR+/HER2+, and HR+ secondary liver tumors, which we used as references for the deconvolution of two liver metastatic MERFISH samples (HER2+ and HR+). In this setting, we observed smaller overall differences in deconvolution performances. Subtype-matched references produced the best results for both samples (HER2+: RMSE *≈* 0.10; HR+: RMSE *≈* 0.17), whereas non-matching references resulted in higher errors (maximum RMSE values of 0.13 and 0.22 for HER2+ and HR+, respectively) (refer **Suppl. Fig, 14**). These findings further supported that deconvolution algorithms are generally robust to reference selection, although subtype-specific matching can still improve their outcomes.

Finally, we assessed the use of large single-cell atlases for deconvolution. We utilized the extensive atlas-scale dataset from Klughammer `et al.` 2024)[27], which included various metastatic samples derived from different patients and tissues (bone, brain, liver, breast etc.). To evaluate technical batch effects derived from multiple distinct patients or data collection protocols on deconvolution performance, we further divided the atlas into two subsets: single-cell and single-nucleus samples. We leveraged three metastatic MERFISH peudospot samples (HR+, HER2+, TNBC+) in this experiment to better represent breast cancer heterogeneity. We observed that the fully integrated Klughammer atlas showed very comparable SSIM (*≈* 0.91) and RMSE (*≈* 0.10) values to the single-nuclei configuration across MERFISH samples. Using the single-cell reference subset however, yielded to worse performances (SSIM *≈* 0.81 and RMSE *≈* 0.17). These findings highlight that Cell2location and RCTD are both effective at mitigating batch effects across samples and even technologies.

### 3.6 Constructing Meta-Cells as the Reference Dataset

In comparison to bulk RNA-seq, scRNA-seq data is sparse. Since each cell are sampled only once, only the mRNA at that specific time point is sequenced. To overcome this, meta-cells are frequently employed in single-cell omics studies. Meta-cells are created by clustering cells of the same type in silico e.g. using the k-nearest-neighbors algorithm [30]. Each cell’s gene expression within the cluster are then compiled to represent a cell across possible growth phases. We created meta-cells from the matched scRNA-seq reference data provided by Janesick `et al.` 2021 [15] and tested to see the impact of deconvolution on the matched Xenium ST data. We found that there was no difference in deconvolution accuracy between using a meta-cell reference (SSIM = 0.80, RMSE = 0.15) and the scRNA-seq reference as is.

## 4 Discussion

This study evalautes the influence of single-cell reference selection and its composition on deconvolution success in breast cancer spatial transcriptomics analysis. Unlike previous benchmarks that mainly compared performance of tools, we focused on the role of reference design, including cell counts and patient or disease subtype matching; large single-cell atlases, and pseudo-bulking methods. Our study was inspired by lessons from genotype imputation and bulk RNA-seq deconvolution, where the diversity and quality of reference panels have been shown to directly shape inference accuracy[37–40]. We performed deconvolution on two Xenium replicates and three MERFISH ST samples in order to generate helpful guidelines for researchers to address reference panel selection. We leveraged different downsampling approaches, pairings of various single samples and large breast and cross-cancer atlases to increase technological and patient-driven heterogeneity in the reference panels.

Several limitations must be addressed in this study. First, our analyses did not include ST samples from every breast cancer sub-type, limiting its generalization. Besides, we focused on using Xenium and MERFISH pseudospots which is only a proxy to mimic Visium-like spots. Different cell-types show various levels of RNA transcription[41–43], making direct associations between cell counts and transcriptional composition of bulk experiments more complicated. On top of that, sequencing-based ST approaches suffer from stronger technical drop-out events than imaging-based ones[44, 45], as well as transcript drift where the RNA molecules from a cell in one spot drift into an adjacent spot[46]; these idiosyncrasies cannot be easily replicated. Additionally, targeted fluorescence based approaches have a strongly limiting gene panel of only a few hundred genes for deconvolution tools to leverage, compared to the unbiased gene capturing of sequencing based approaches like Visium(HD).

Furthermore, the process of segmenting and annotating cells in imaging-based ST technologies presents challenges. These methods capture RNA transcripts efficiently but are still found lacking in comparison to contemporary scRNA-seq protocols. Machine-learning-based cell segmentation is still not fully reliable with challenges including misassigning reads to neighboring cells, detecting cells of different sizes and compositions, and correctly identifying overlapping cells. [46, 47] Consequently, ground truth cell-type annotations from Xenium and MERFISH potentially skew deconvolution performance evaluations.

Additional errors in cell-type subtyping for scRNA-seq references, particularly in larger atlas-scale datasets, can significantly affect deconvolution results. Discrepancies in cell-type annotations or annotation level between different studies, as well as imperfect reliability of annotation tools, contribute to inconsistencies in the labeling process.

Evaluation of deconvolution results revealed important differences across performance metrics. Global measures such as RMSE and SSIM were highly consistent, whereas cell-type–specific evaluations captured complementary aspects: RMSE penalized large point-wise deviations, while JSD and PCC highlighted structural agreement. These discrepancies uncovered a systematic underestimation of cell types strongly enriched in specific spots, such as tumor or stromal cells in primary breast cancer deconvolution. The presence of a few highly abundant cell types within spatial spots has already been recognized as a difficult scenario for these algorithms by Sang-Aram `et al.`[23], and our findings confirm this limitation. Such systematic underestimation is particularly concerning, as it can artificially inflate inferred immune contributions and lead to potential clinical misinterpretations such as false-positive aggregation of immune cells in tumors.

By contrast, cell types with moderate abundance were consistently well recovered, achieving the highest performances across metrics. But detecting rare populations also posed a different challenge: PVL cells in primary breast cancer and B cells in metastatic samples were not overestimated but their spatial patterns were hardly detected. This aligns with previous reports highlighting the inherent difficulty of resolving rare cell types via deconvolution[23, 43]. Together, these findings underscore that current deconvolution methods still struggle with both dominant and rare cell types in spot-based ST data. This study showed that this result did not markedly improve with a variety of cell proportions of these rare cells within the reference, pointing to either a flaw in how current state-of-the-art tools interpret data from these cell types, or another unknown technical bias. Consequently, inferred cell-type maps must be interpreted with caution, particularly when translated into biological or clinical conclusions.

Surprisingly, we observed that the number of cells and cell-type distribution within the single-cell references did not play an important role in guiding cellular deconvolution as long as the input reference is of high-quality. RCTD requires at least 25 cells per cell-type while Cell2location could possibly achieve reliable deconvolution results with even less cells. This means that researchers are free of the large computational and monetary burden of collecting vast reference datasets, as long as every cell-type is represented with a small amount of cells. Thus, for breast cancer and perhaps most other tissue types, even small scRNA-seq references of around 500 cells should be sufficient for accurate deconvolution.

We further showed that both RCTD and Cell2location achieved higher overall deconvolution accuracy when distinguishing broader-grained cell-type annotations (8 types) compared to finer categories (17 types), likely reflecting the reduced complexity of the assignment task at coarser resolutions. Nonetheless, cell-type–specific performances varied: some populations, such as T cells, benefited markedly from subtyping, whereas others, including cancer cells, showed opposite effects. We therefore recommend performing deconvolution at both annotation levels and interpreting sub–type-specific differences with caution.

Pairing primary breast cancer ST samples with different scRNA-seq references showed that matched single-cell datasets provide only modest performance gains but yield more stable deconvolution results, with Cell2location exhibiting a stronger dependence on matched references than RCTD. In concordance, varying the degree of subtype matching for metastatic MERFISH samples again yielded more stable results when considering the underlying molecular classifications for reference selection. Nonetheless, both tools demonstrated that they can robustly capture gene expression patterns regardless of breast cancer subtype or collection protocol.

Finally, we evaluated differing technical makeups of the reference including large single-cell datasets. Atlases diminish the necessity of feature engineering and increasing gene expression heterogeneity by encompassing a wide diversity of patients with different clinical backgrounds[31]. In the primary breast cancer samples, we demonstrated that large breast cancer atlases such as from Wu `et al.` 2021[33] can lead to deconvolution performances almost on par with matching references. Testing the limits of deconvolution, we also evaluated the usage of cross-cancer datasets. These results demonstrated that even some non-breast cancer panels were able to provide useful data for deconvolution (e.g. Ovarian cancer atlas by Olbrecht `et al.`[35] for RCTD), likely due to conserved malignant expression signatures. However, we show that performance strongly varied between computational tools and reference combinations. We therefore encourage the leveraging of the vast resources of publicly available single-cell datasets as reference panels for deconvolution, but stress caution that these atlases do not guarantee success. This advice might be especially important for researchers that cannot perform additional single-cell experiments on top of their ST studies, through lack of funding or sample availability. Resources like the *Human Cell Atlas*[48], *Curated Cancer Cell Atlas* [49] *Tabula sapiens*[50], or *Tabula muris*[51] for mouse offer robust, well-annotated and extensive datasets for researchers to utilize.

On the other hand, pseudo-bulk meta-cells offered an opportunity to evaluate how density of the data affected deconvolution success. There was essentially no difference between the use of the scRNA-seq reference or a meta-cell reference built on the same matched sample. Even though the meta-cell reference contained fewer cells, there was no significant improvement in computational time for deconvolution. In any case, any improvement was offset by the time taken to compute the meta-cells from the raw data input.

Looking ahead, advances in high-resolution spatial transcriptomics protocols such as Stereo-seq and VisiumHD promise to achieve subcellular resolution while still providing an unbiased capture of transcripts across the tissue[52, 53]. These improvements may reduce the need for traditional spot-level deconvolution, enabling a shift toward approaches based on precise cell segmentation and direct assignment of transcripts to individual cells[54, 55]. Nevertheless, given current limitations in capture efficiency and segmentation accuracy, computational deconvolution will remain an essential tool for linking single-cell profiles with spatial context and extracting biologically meaningful insights for the foreseeable future.

## 5 Conclusions

When performing spatial transcriptomics deconvolution, we advise using matched single-cell references whenever possible. However, samples from the same breast cancer subtype or even other patient samples from different subtypes can also yield stable and reliable deconvolution results, highlighting the robustness of algorithms such as RCTD and Cell2location with respect to reference choice. Large breast cancer atlases, which capture greater heterogeneity, can serve as effective reference datasets and often perform nearly as well as matched samples. The accuracy of deconvolution varies across cell types depending on their abundance in the tissue, with highly abundant or very rare cell types posing particular challenges. Subtyping can improve deconvolution for certain heterogeneous populations, such as myeloid cells or T cells, but does not necessarily enhance overall performance. Therefore, deconvolution results should always be interpreted with care, considering both the characteristics of the reference dataset and the cell-type-specific limitations.

## 6 List of abbreviations

ATC: Anaplastic Thyroid Cancer
BC: Breast Cancer
C2L: Cell2location
CAFs: Cancer-Associated Fibroblasts
DCIS: Ductal Carcinoma In Situ
ER: Estrogen Receptor
HER2: Human Epidermal Growth Factor 2
HGSTOC: High-Grade Serous Tubo-Ovarian cancer
HR: Hormonal Receptor
IHC: Immunohistochemistry
IDC: Invasive Ductal Carcinoma
ILC: Invasive Lobular Carcinoma
JSD: Jensen-Shannon Divergence
NGS: Next-Generation Sequencing
MBC: Metastatic Breast Cancer
PCC: Pearson Correlation Coefficient
PR: Progesterone Receptor
PVL: Perivascular-like Cells
RMSE: Root Mean Squared Error
RCTD: Robust Cell Type Decomposition
scRNA-seq: single-cell RNA-sequencing
SSIM: Structural Similarity Index
ST: Spatial Transcriptomics
TAM: Tumor-Associated Macrophages
TIL: Tumor Infiltrating Lymphocytes
TME: Tumor Micro-Environment
TNBC: Triple Negative Breast Cancer

## 7 Declarations

### 7.1 Availability of data and materials

All datasets used in this study are publicly accessible. Single-cell and Xenium datasets from Janesick `et al.` were obtained from the Gene Expression Omnibus (GE0) database under *GSE243280* and cell-type annotations from the attached *10x Genomics entry*. Processed scRNA-seq data from Wu `et al.` was downloaded from GEO under the accession number *GSE176078*. Curated single-cell atlases from Gao[34], Olbrecht[35] and Pelka[36] were downloaded by the *Curated Cancer Cell Atlas* platform from the Weizmann Institute. Data bundles containing single-cell/nuclei and MERFISH samples from Klughammer were retrieved from the *Single-Cell Portal*

The full analysis workflow, including code and documentation, is available on GitLab (https://gitlab.com/daub-lab/stdeconvref).

### 7.4 Author’s contribution

Conceptualization: S.A., S.W., C.D. Data curation: S.A. Formal analysis: S.A., S.W. Methodology: S.A., S.W. Writing – original draft: S.A. Writing – review & editing: S.A., S.W., C.D.

### 7.5 Acknowledgements

We thank the members of the Daub Lab, namely Rasha Fahad, Gabriele Gambero, Faezeh Mottaghitalab, Işıl Takan and Rijja Hussain Bokhari for helpful discussions and feedback, as well as the members of the Spatial Transcriptomics for Breast Cancer group, including Magnus Boman, Alexis Ioannou, Maria Arceo and Cevi Bainton for their valuable input and collaborative exchange. We are also grateful to Stephan Ossowski and Manfred Claassen at the University of Tübingen for their guidance and constructive suggestions. We thank the Center for Bioinformatics and Biostatistics (CBB) at Karolinska Institutet for compute resources.

### 7.6 AI usage statement

Portions of the manuscript were edited with the assistance of AI-based tools (OpenAI ChatGPT, GPT-4). The authors reviewed and verified all content. AI-assisted code generation (OpenAI ChatGPT, GPT-4) was used for drafting data analysis scripts. All code was reviewed and validated by the authors.

