## Supplementary File for "Impact of single-cell RNA reference selection for the deconvolution of breast cancer spatial transcriptomics datasets"

Table S1: Summary of spatial transcriptomics datasets used.

**Abbreviations:** BC = breast cancer; IDC = invasive ductal carcinoma; ILC = invasive lobular carcinoma; DCIS = ductal carcinoma in situ; MBC = metastatic breast cancer; HR = hormonal receptor; HER2 = human epidermal growth factor receptor.

| **Study** | **Sample** | **Technology** | **Tissue** | **Histology** | **Clinical** | **Pseudospots** |
|---|---|---|---|---|---|---|
| Janessick `et al.` 2023 | Sample 1 R1 | Xenium | BC | DCIS | HR+/HER2+ | 12,710 |
| Janessick `et al.` 2023 | Sample 1 R2 | Xenium | BC | DCIS | HR+/HER2+ | 11,373 |
| Klughammer `et al.` 2024 | 313-932 | MERFISH | Liver Mets | MBC | HER2+ | 1,726 |
| Klughammer `et al.` 2024 | 944-7479 | MERFISH | Liver Mets | MBC | HR+ | 1,110 |
| Klughammer `et al.` 2024 | 917-4531 | MERFISH | Liver Mets | MBC | TNBC | 1,673 |

Table S2: Summary of single-cell reference datasets used.

| **Study** | **Sample** | **Technology** | **Tissue** | **Histology** | **Clinical** | **Cells** |
|---|---|---|---|---|---|---|
| Wu et al. 2021 | CID4290 | scRNA-seq | BC | IDC | HR+ | 5,789 |
| Wu et al. 2021 | CID4465 | scRNA-seq | BC | IDC | HR+ | 1,564 |
| Wu et al. 2021 | CID44971 | scRNA-seq | BC | IDC | TNBC | 7,986 |
| Wu et al. 2021 | CID4535 | scRNA-seq | BC | ILC | TNBC | 3,961 |
| Janessick et al. 2023 | Sample 1 | scRNA-seq | BC | DCIS | HR+/HER2+ | 21,778 |
| Wu et al. 2021 | Atlas | scRNA-seq | BC | IDC, ILC | - | 100,064 |
| Gao et al. 2021 | Atlas | scRNA-seq | BC, TC | DCIS, TNBC, ATC | - | 24,704 |
| Olbrecht et al. 2021 | Atlas | scRNA-seq | OV | HGSTOC | - | 12,411 |
| Pelka et al. 2021 | Atlas | scRNA-seq | CRC | AdCa | - | 313,220 |
| Klughammer et al. 2024 | 313-932 | scRNA-seq | Liver Mets | MBC | HER2+ | 12,468 |
| Klughammer et al. 2024 | Merged | scRNA-seq | Liver Mets | MBC | HER2+ | 12,492 |
| Klughammer et al. 2024 | Merged | scRNA-seq | Liver Mets | MBC | HR+/HER2+ | 13,167 |
| Klughammer et al. 2024 | Merged | scRNA-seq | Liver Mets | MBC | HR+ | 15,086 |
| Klughammer et al. 2024 | Atlas | scRNA-seq | MBC | MBC | - | 33,708 |
| Klughammer et al. 2024 | Atlas | snRNA-seq | MBC | MBC | - | 72,743 |
| Klughammer et al. 2024 | Atlas | Mixed | MBC | MBC | - | 106,451 |

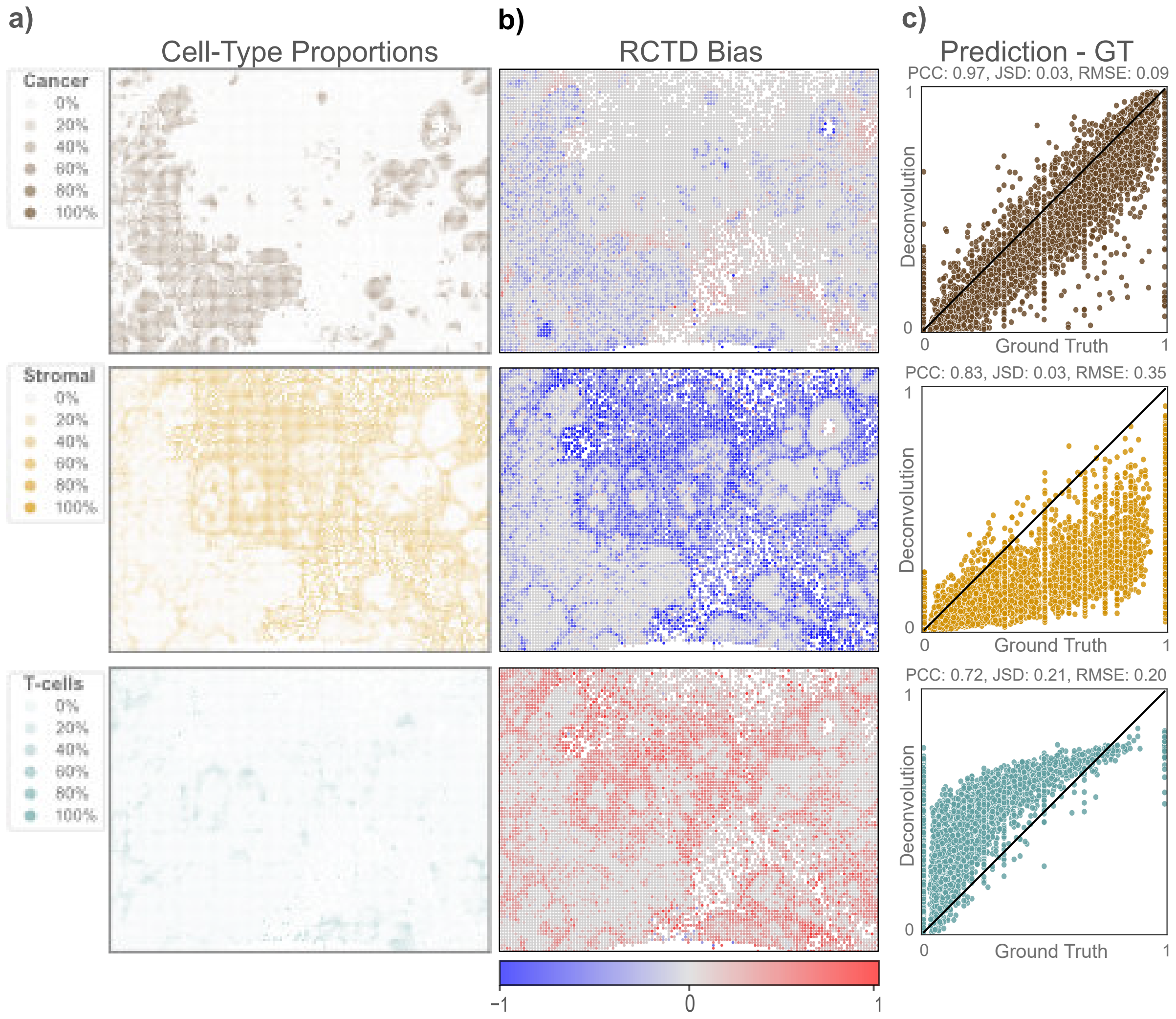


Figure S1: Deconvolution analysis results for three key cell types: cancer cells, stromal cells, and T-cells (analog to Fig. 1). **a)** Cell-type proportions per spatial spot based on segmented Xenium cell annotations. **b)** Difference between deconvolved proportions and ground truth for each spot, with overestimation indicated by red and underestimation by blue. **c)** Correlation plots comparing predicted and ground truth proportions for each spot, with the diagonal line representing perfect prediction. Performance metrics, including Pearson Correlation Coefficient (PCC), Jensen-Shannon Divergence (JSD), and Root Mean Squared Error (RMSE), are provided to quantify the accuracy of the predictions.

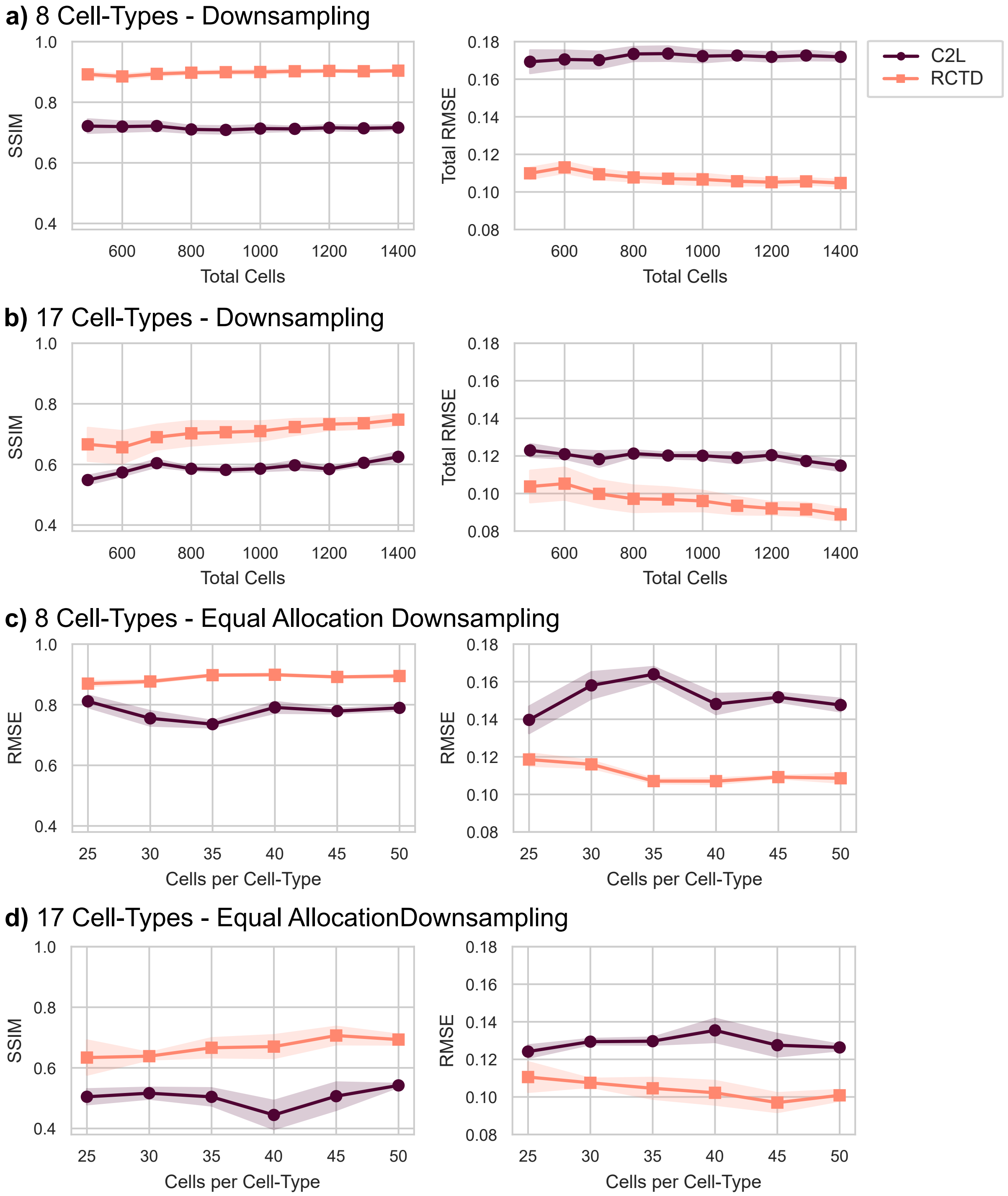


Figure S2: Global deconvolution performance metrics (SSIM, RMSE) for stratified and equal allocation downsampling results generated by RCTD and Cell2location . Xenium replicate 1 from Janesick`et al.` was deconvolved by using its matching single-cell reference.

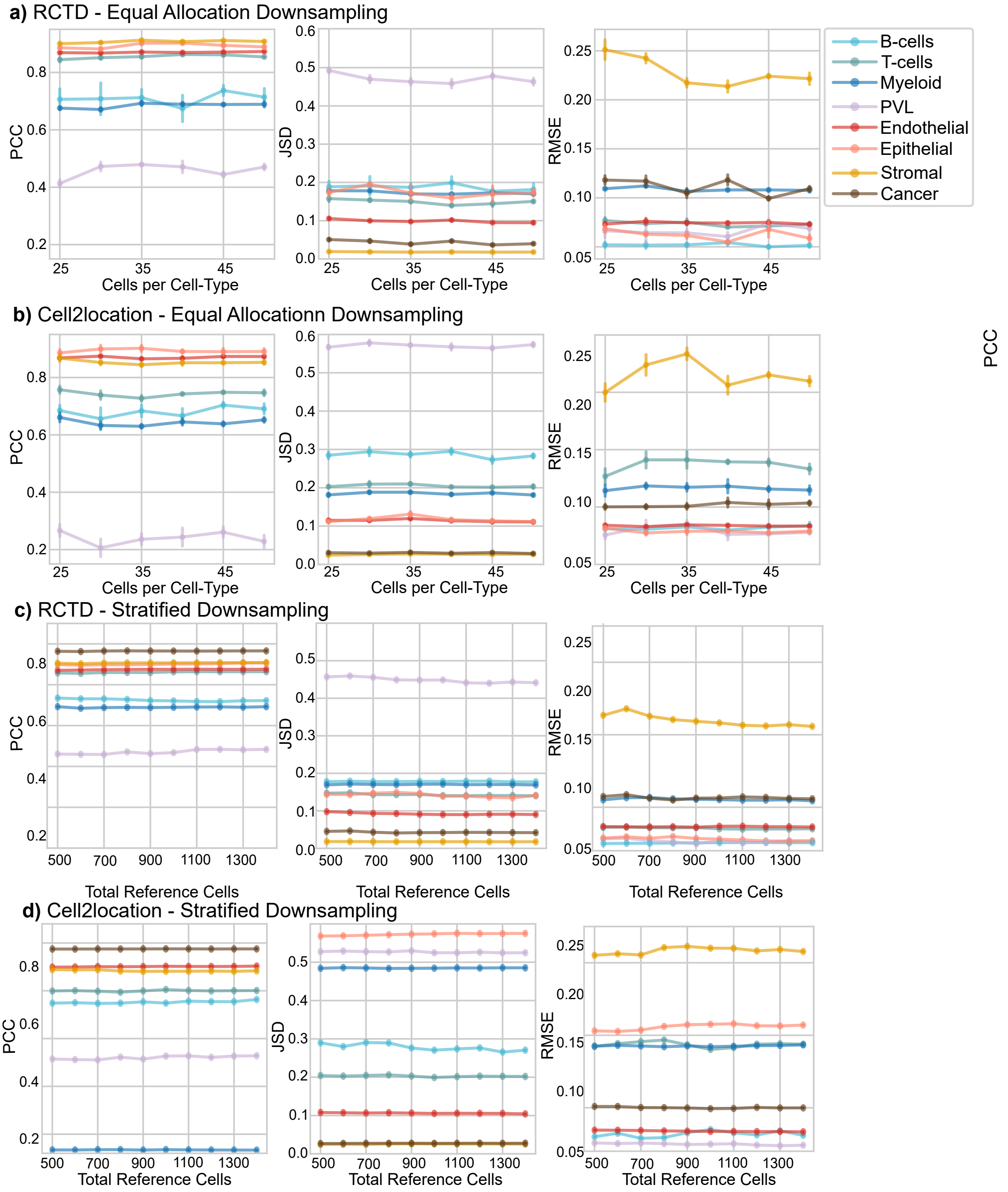


Figure S3: Cell-type-specific performance metrics (PCC, JSD and RMSE) for stratified and equal allocation downsampling results generated by RCTD and Cell2location. Xenium replicate 1 from Janesick`et al.` was deconvolved by using its matching single-cell reference.

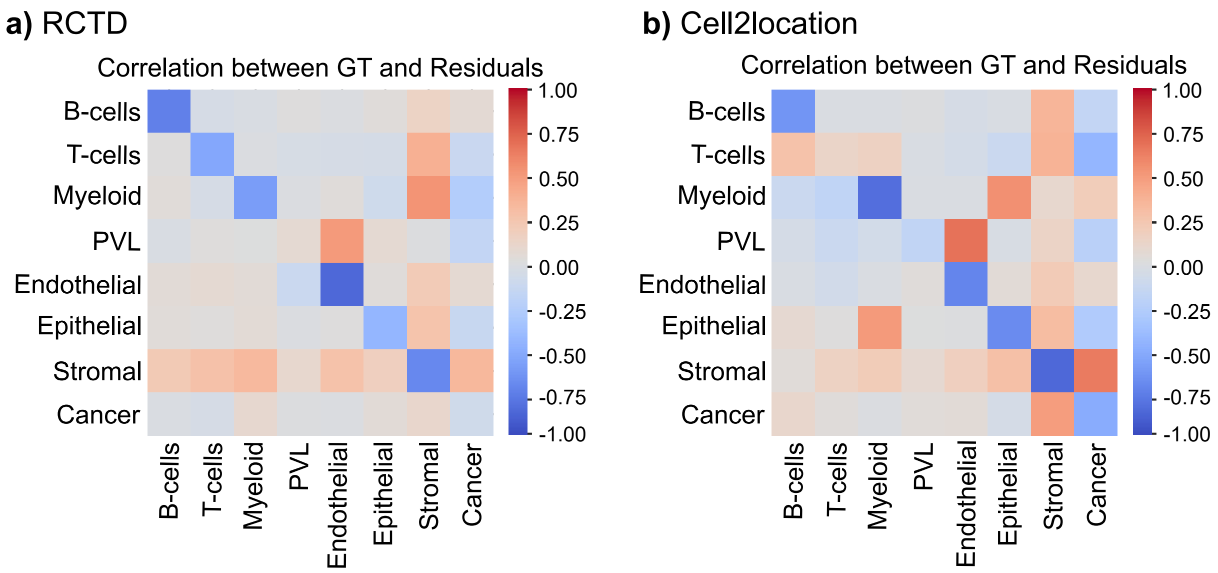


Figure S4: Correlation between ground truth proportions and deconvolution residuals for **a** RCTD and **b** Cell2location. Residuals were calculated by subtracting the ground truth cell-type proportion from its corresponding prediction value for each spot.

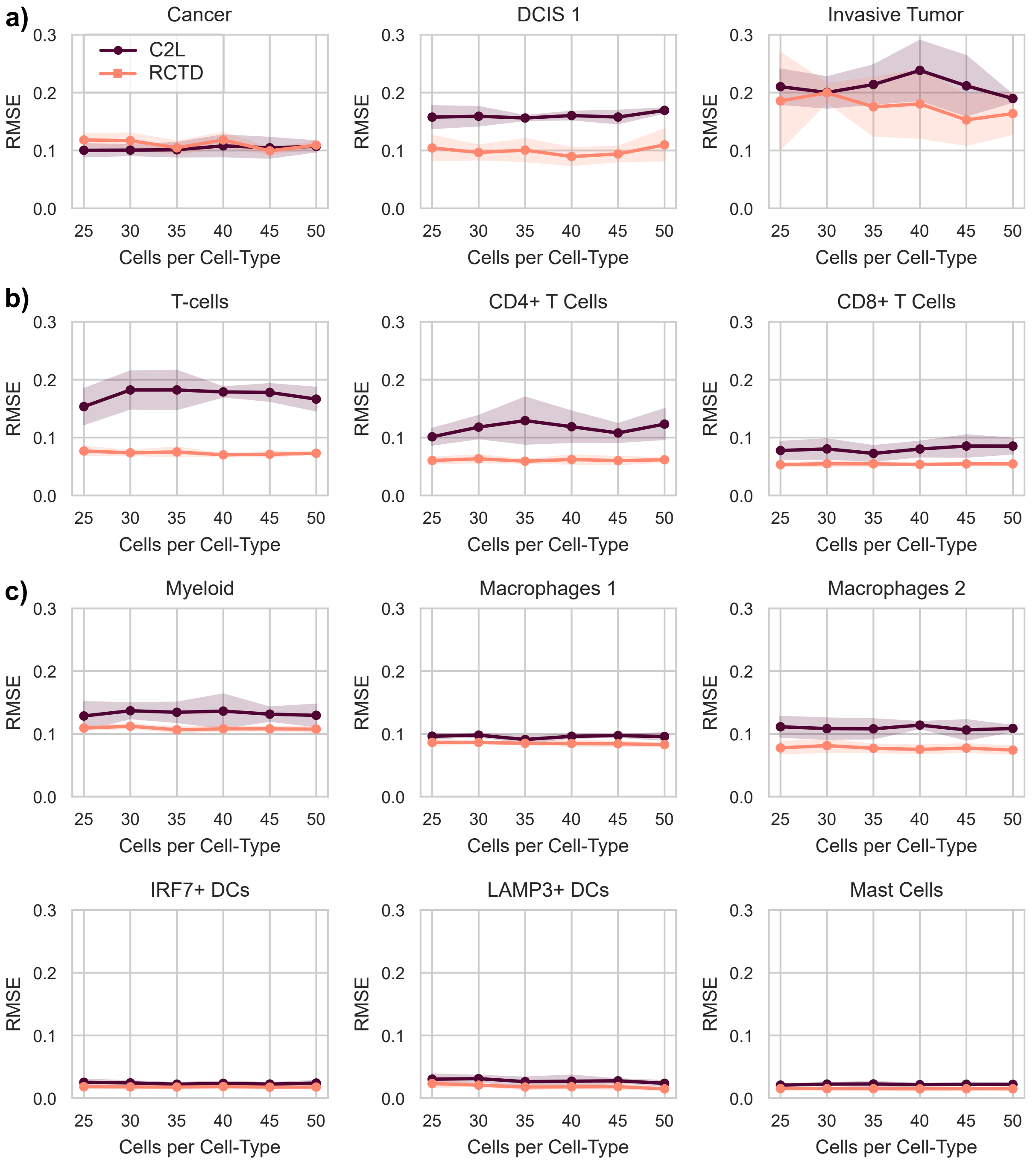


Figure S5: Equal allocation downsampling results for broad -and finer-grained cell-type deconvolution results. **a)** Deconvolution performances across cancer cells and their ductal carcinoma in situ 1 (DCIS 1) and invasive tumor subtypes. **b)** Comparison between RMSE values for T-cells and subtypes CD4+, CD8+ T-cells. **c)** Myeloid RMSE performances across macrophages (Macrophages 1, Macrophages 2), dendritic cells (IRF7+ DCs, LAMP3+ DCs) and mast cells

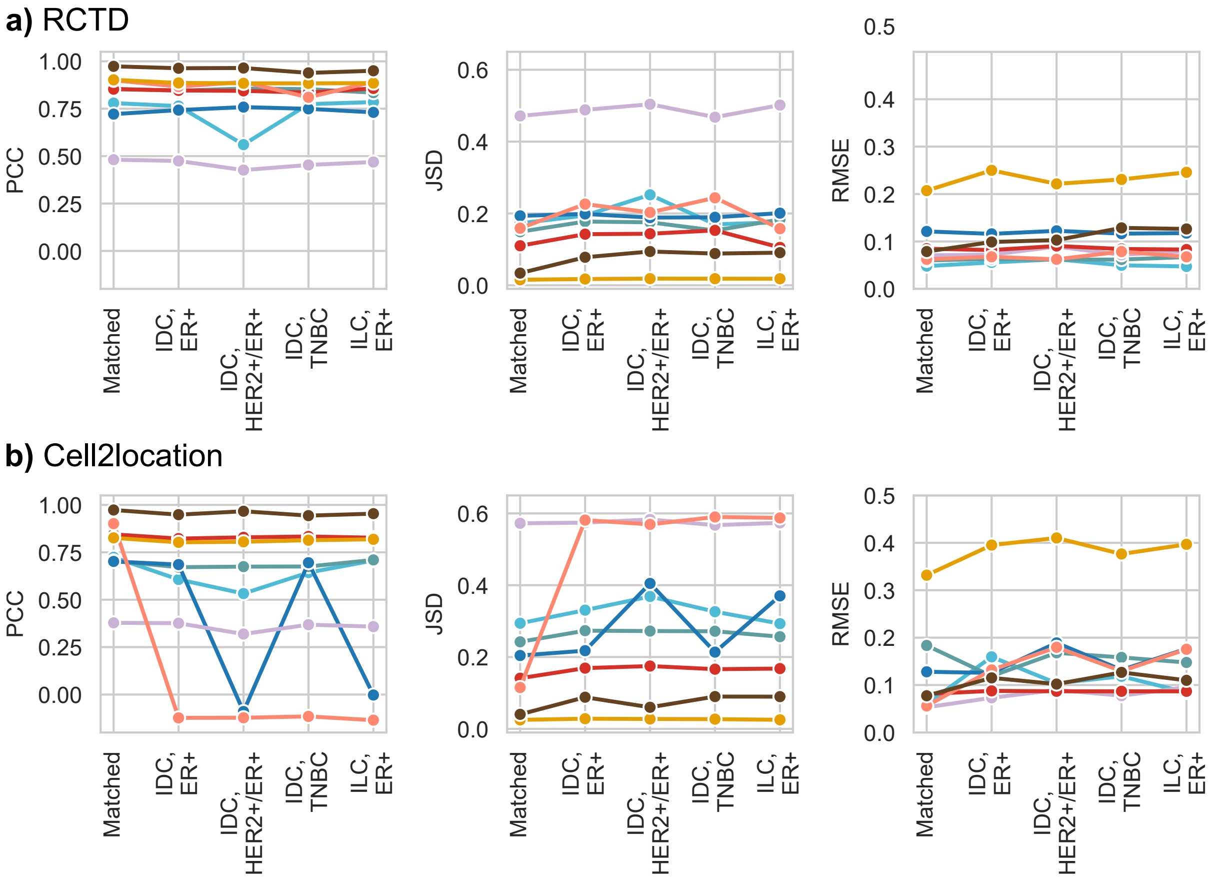


Figure S6: Cell-type specific performances across different ST-sample pairings for **a)** RCTD and **b)** **Cell2location** in Xenium pseudospots.

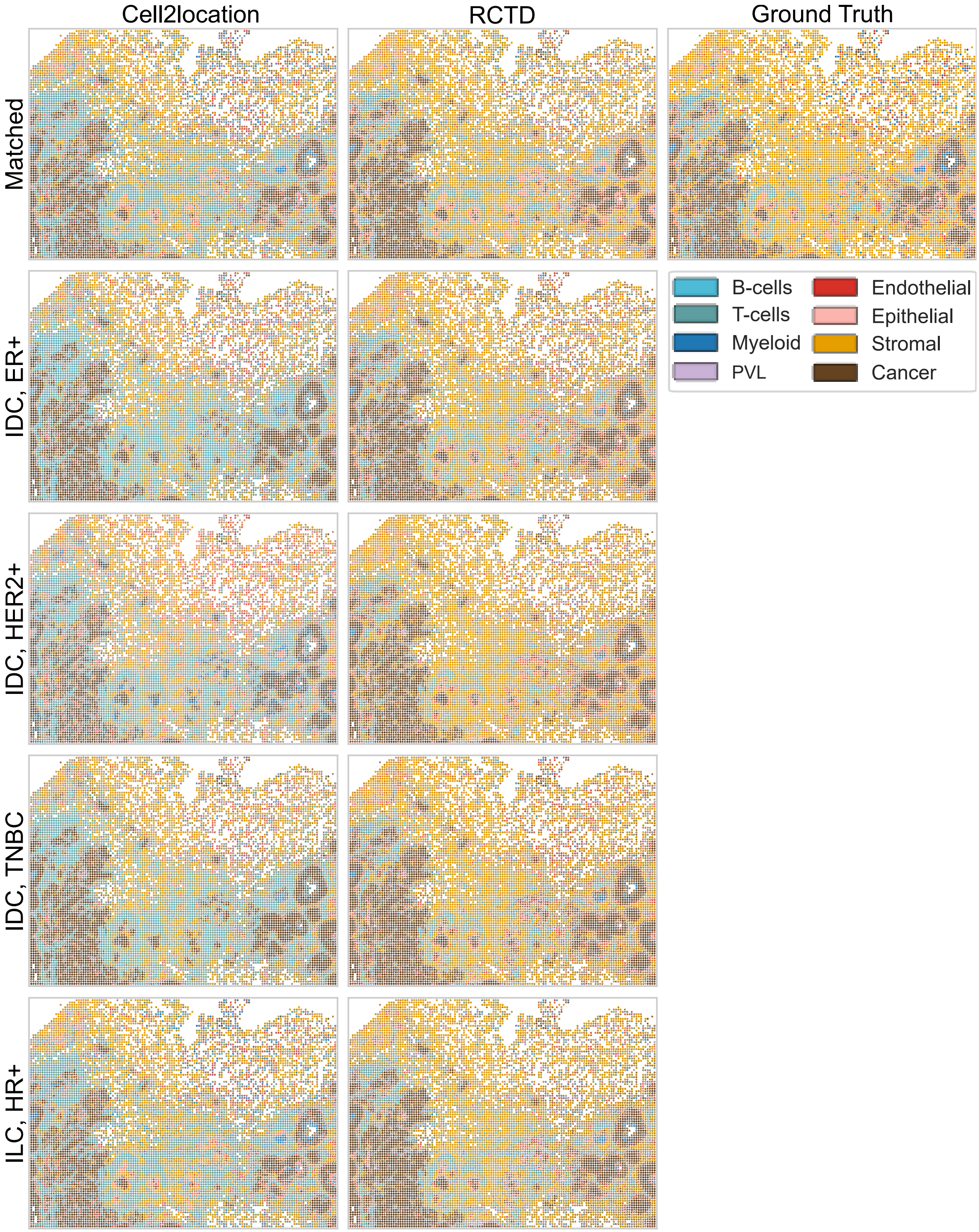


Figure S7: Deconvolution results illustrated as spatial pie plots for different single-cell sample reference pairings. Comparison to the underlying tissue architecture of Xenium replicate 2 from Janesick`et al`.

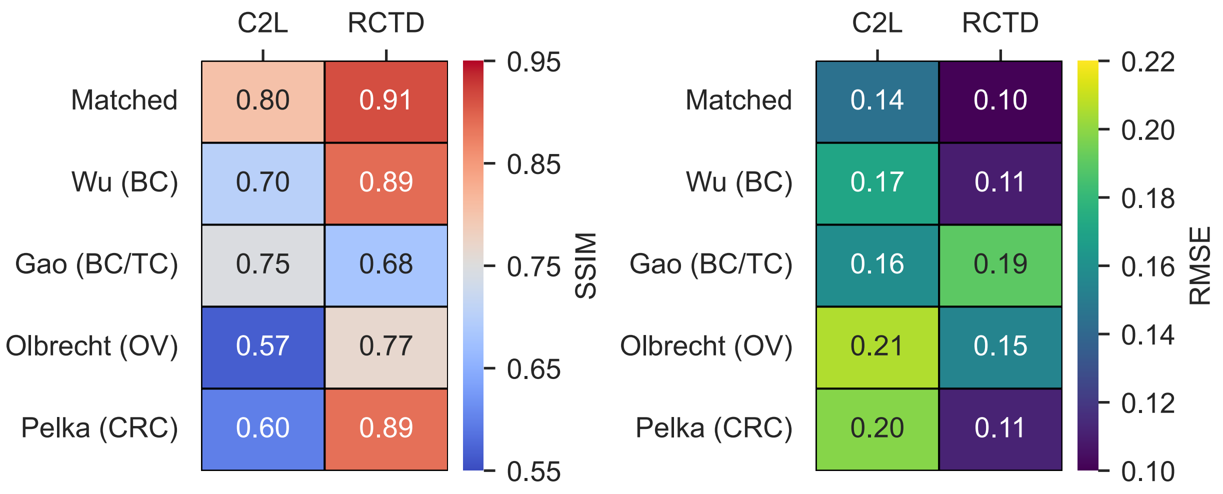


Figure S8: Global deconvolution performances of RCTD and Cell2location by using different single-cell reference atlase. Shown values are the mean performances across both Xenium replicates 1 and 2.

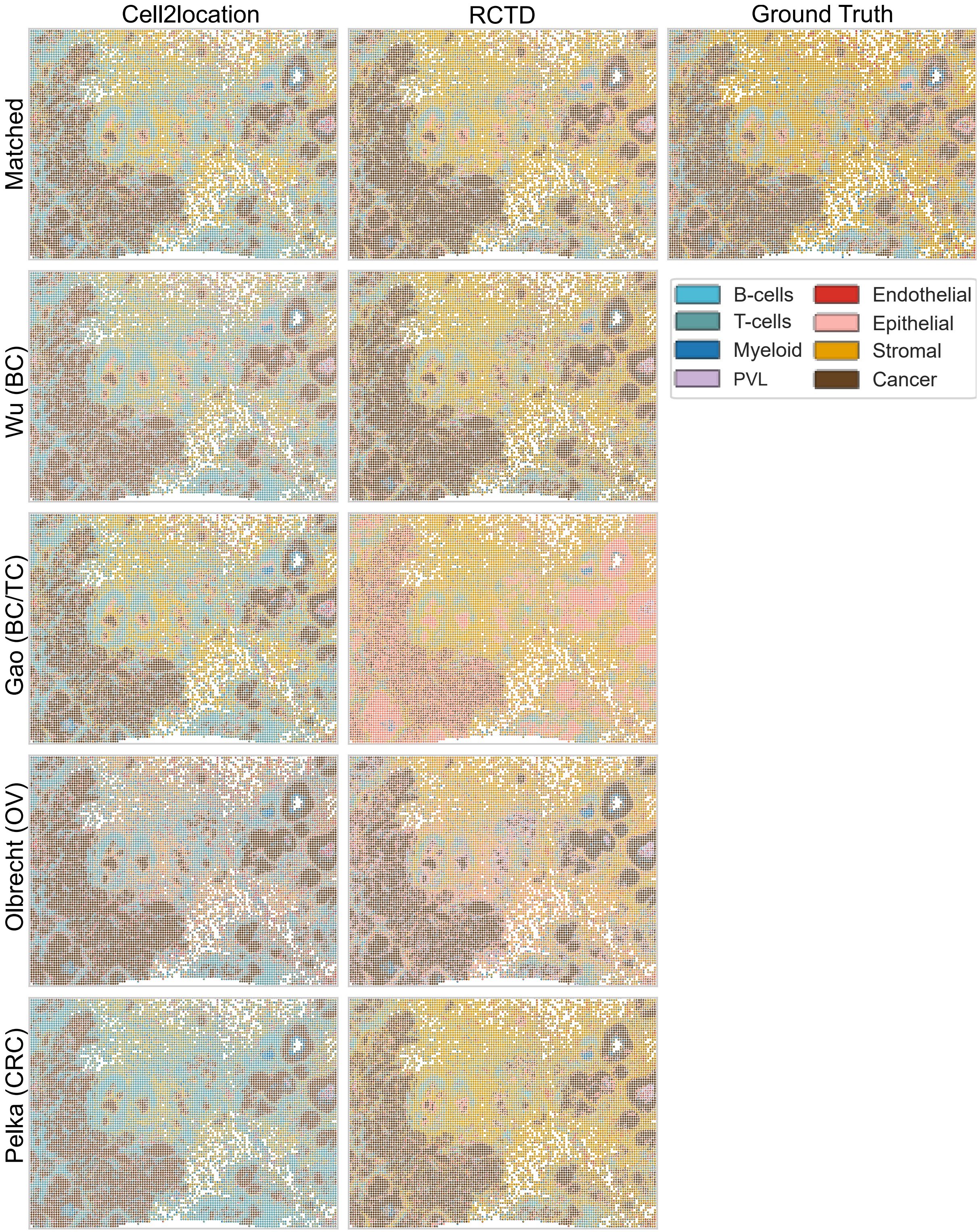


Figure S9: Deconvolution results illustrated as spatial pie plots for different single-cell reference atlases. Comparison to the underlying tissue architecture of Xenium replicate 1 from Janesick`et al`.

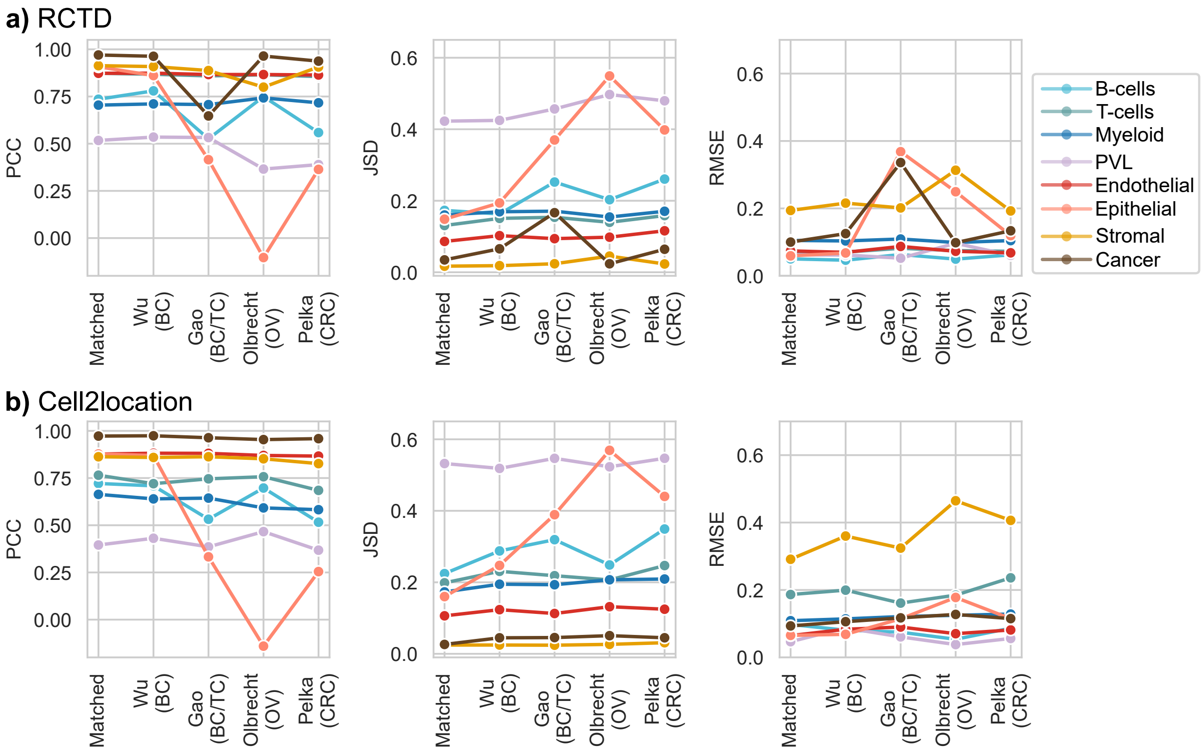


Figure S10: Cell-type specific performances across different ST-sample atlas pairins for **a)** RCTD and **b) Cell2location** in Xenium pseudospots.

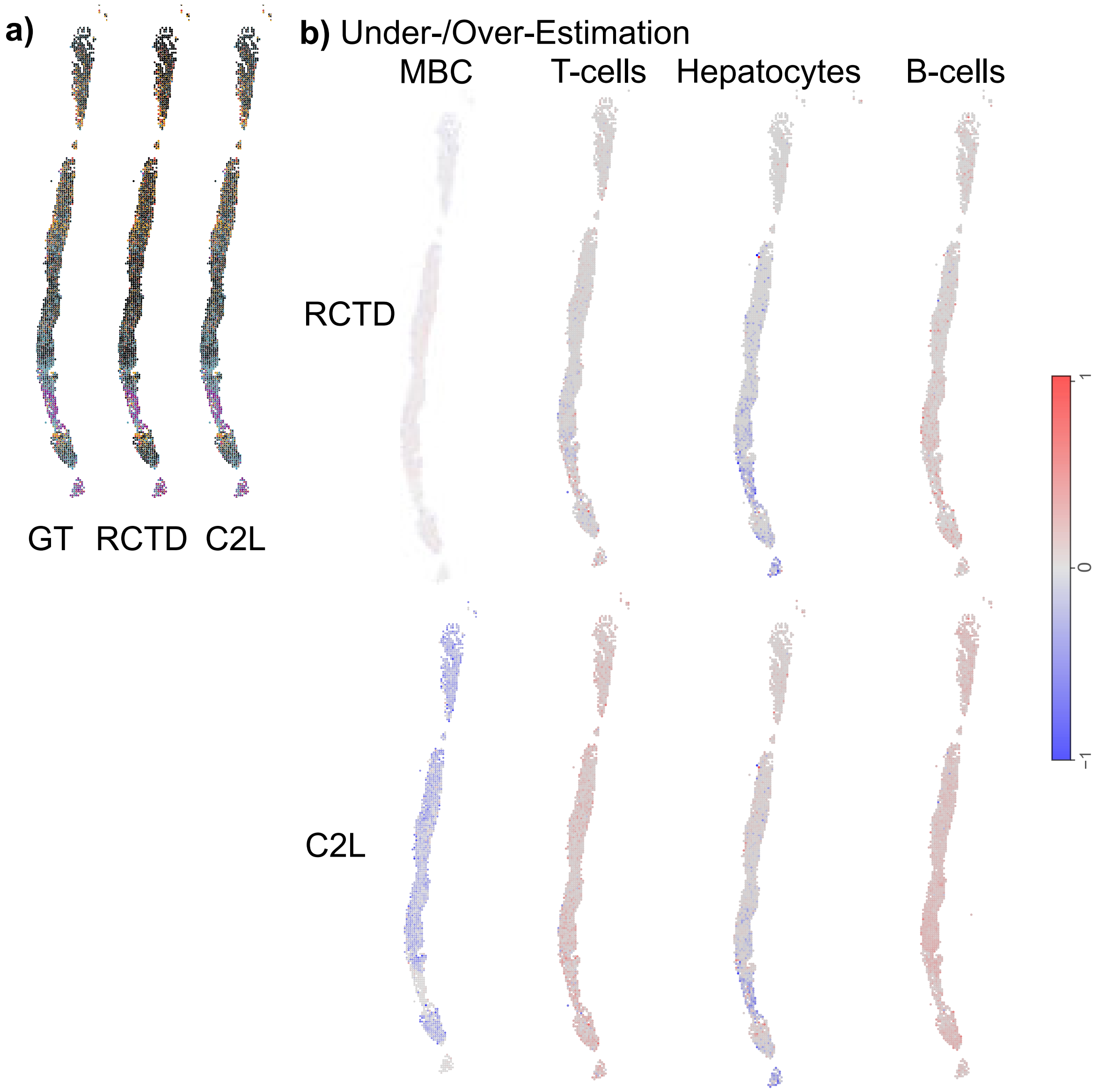


Figure S11: Visualization of MERFISH deconvolution while using a matching single-cell reference. **a)** Inferred cell-type compositions illustrated as spatial pie plots for each spot. **b)** Analysis of cell-type specific under/over-estimations by calculating the difference from predicted to ground truth proportions.

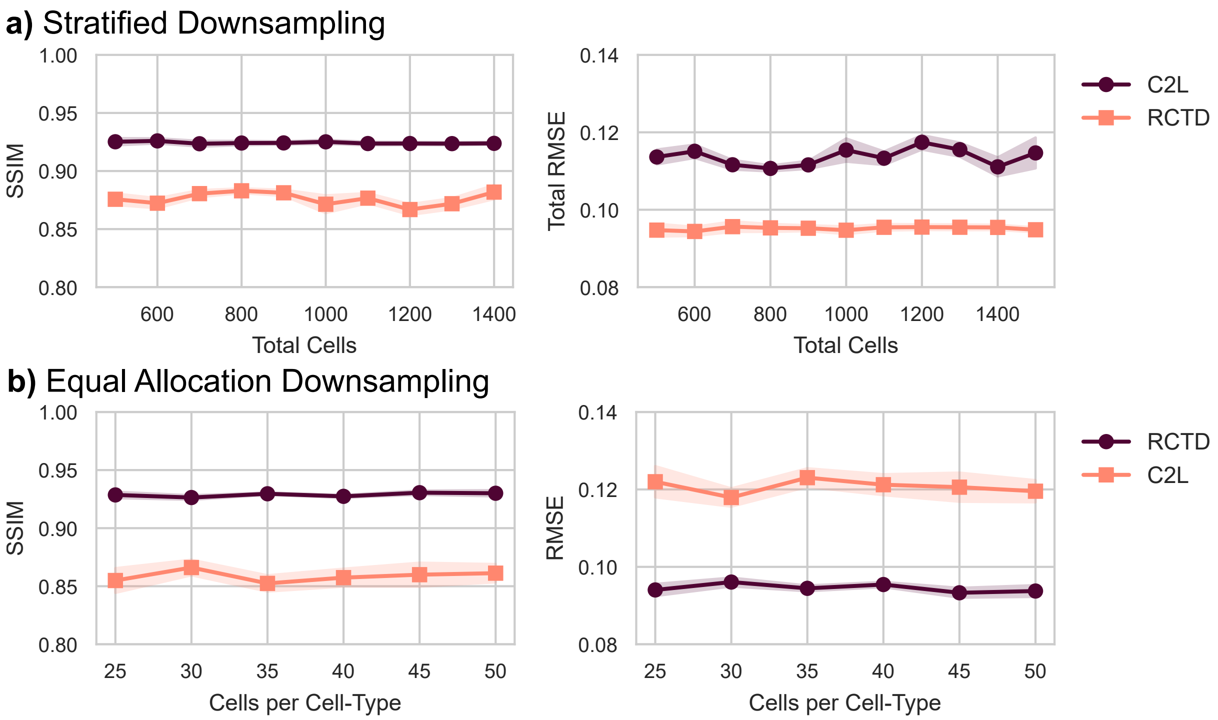


Figure S12: Global deconvolution performance metrics (SSIM, RMSE) for stratified and equal allocation downsampling results generated by RCTD and Cell2location. MERFISH sample 313-932 from Klughammer`et al.` was deconvolved by using its matching single-cell reference.

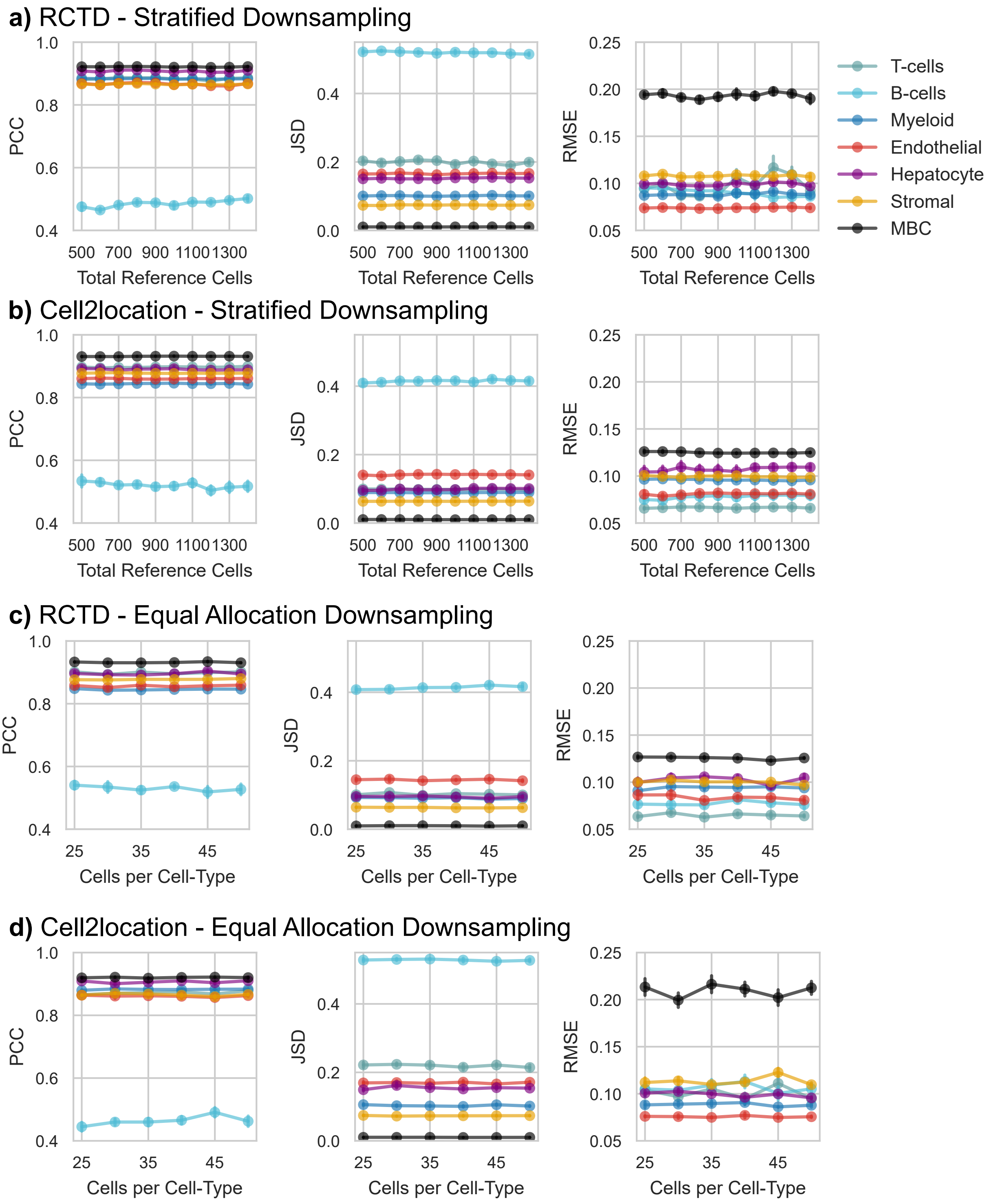


Figure S13: Cell-type-specific performance metrics (PCC, JSD and RMSE) for stratified and equal allocation downsampling results generated by RCTD and Cell2location. MERFISH sample 313-932 from Klughammer`et al.` was deconvolved by using its matching single-cell reference.

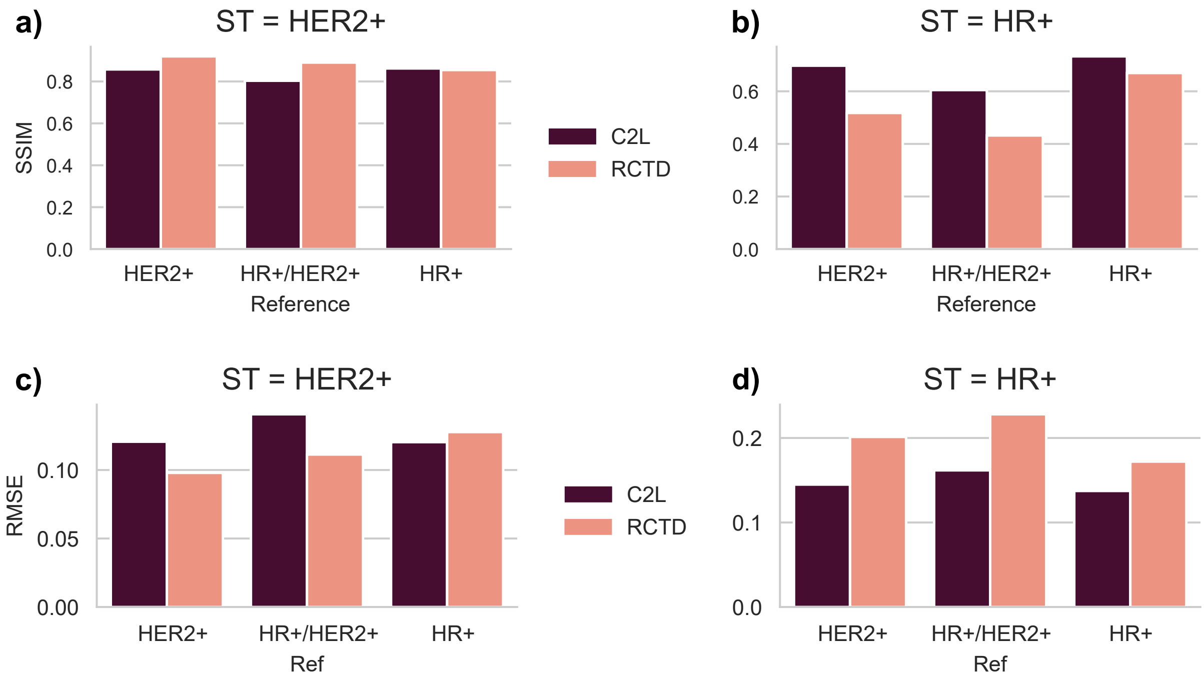


Figure S14: Global deconvolution performances for Cell2location and RCTD across different MERFISH-reference pairings. **a)** and **b)** demonstrate SSIM evaluations for the HR+ and HER2+ MERFISH sample respectively. **c)** and **d)** show RMSE performances.

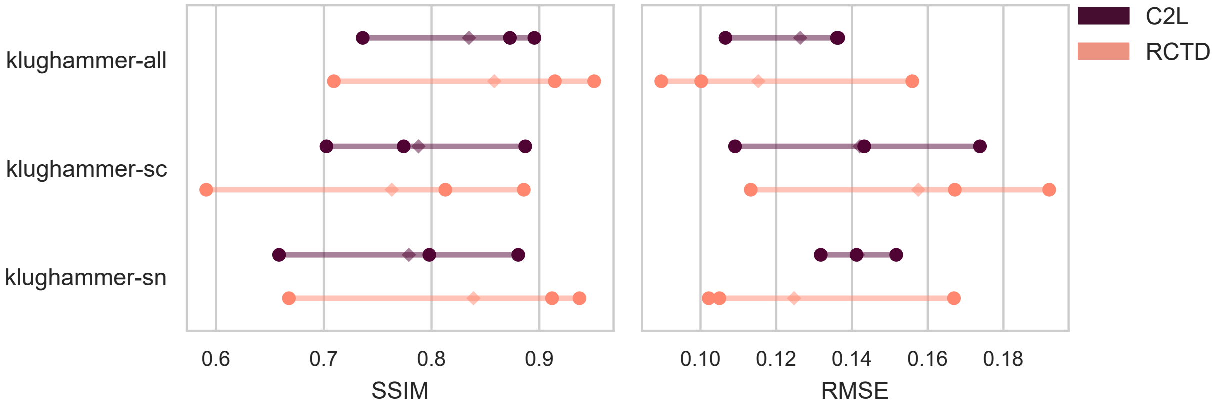


Figure S15: Global deconvolution performances for Cell2location and RCTD across different MERFISH-atlas pairings. Three MERFISH samples (HR+, HER2+, TNBC) were deconvolved via the klughammer single-cell atlas in three configurations: all samples (klughammer-all), single-cell only (klughammer-sc) and single-nuclei only (klughammer-sn).
